## Supplementary material for "Visual working memories are abstractions of percepts": Supp Mats

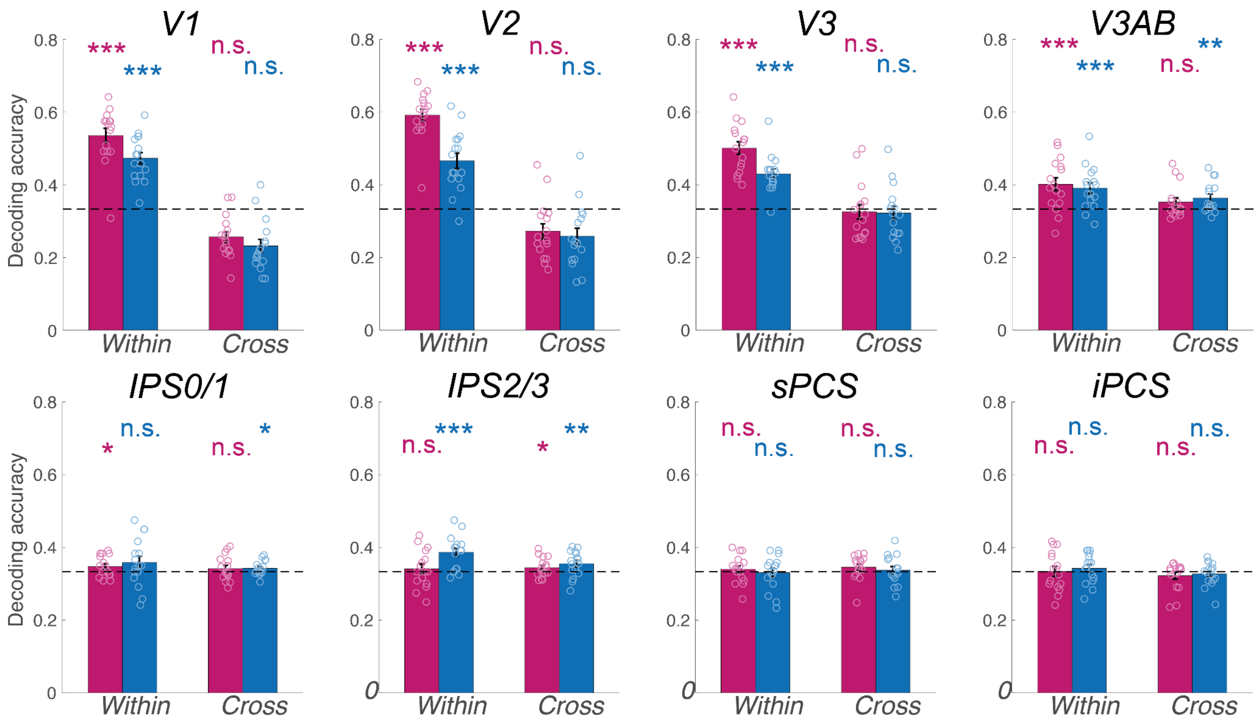


**Figure S1**. Within and cross-modulator decoding results by using the stimulus period of the perceptual control task. *$p$<.05, **$p$<.01, ***$p$<.001, n.s. Not significant. Error bars represent ±1 SEM. Small circles for each bar represent individual data. Dashed horizontal line denotes theoretical chance level (1/3), but results are based on non-parametric permutation tests.

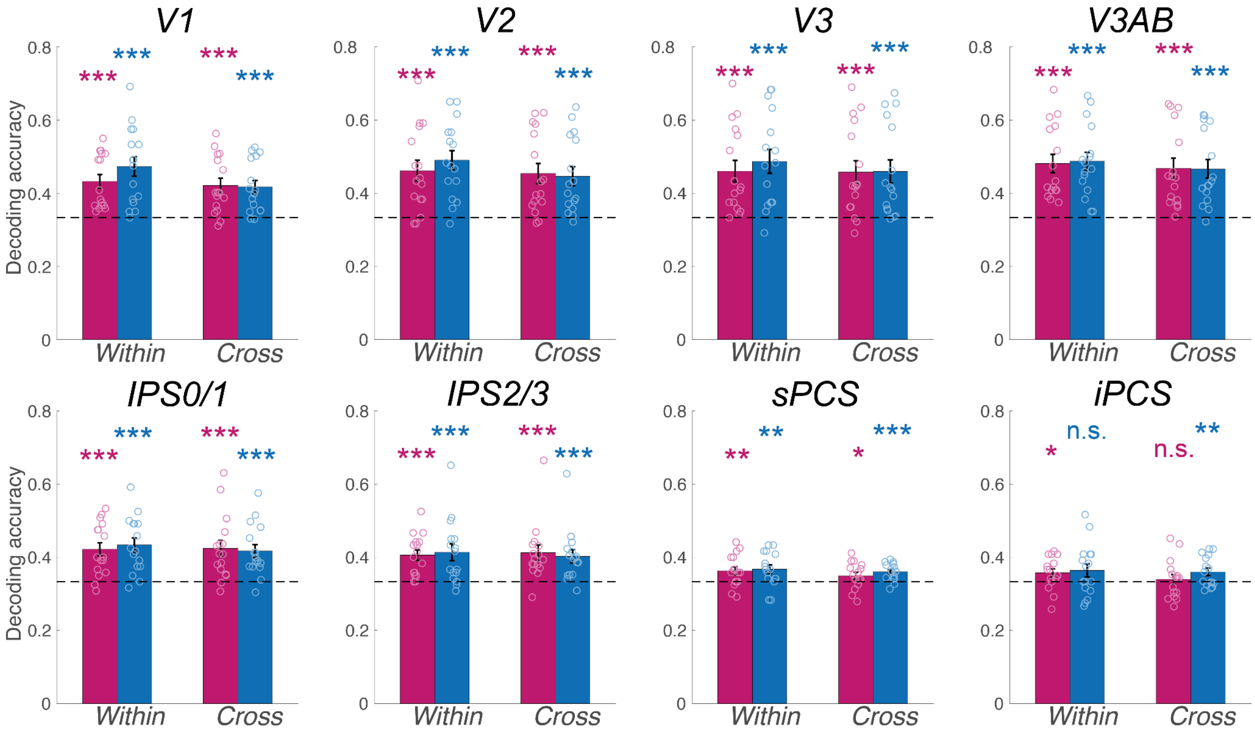


**Figure S2.** Within and cross-modulator decoding results by using the late delay period of the WM task. *$p$<.05, **$p$<.01, ***$p$<.001, n.s. Not significant. Error bars represent ±1 SEM. Small circles for each bar represent individual data. Dashed horizontal line denotes theoretical chance level (1/3), but results are based on non-parametric permutation tests.

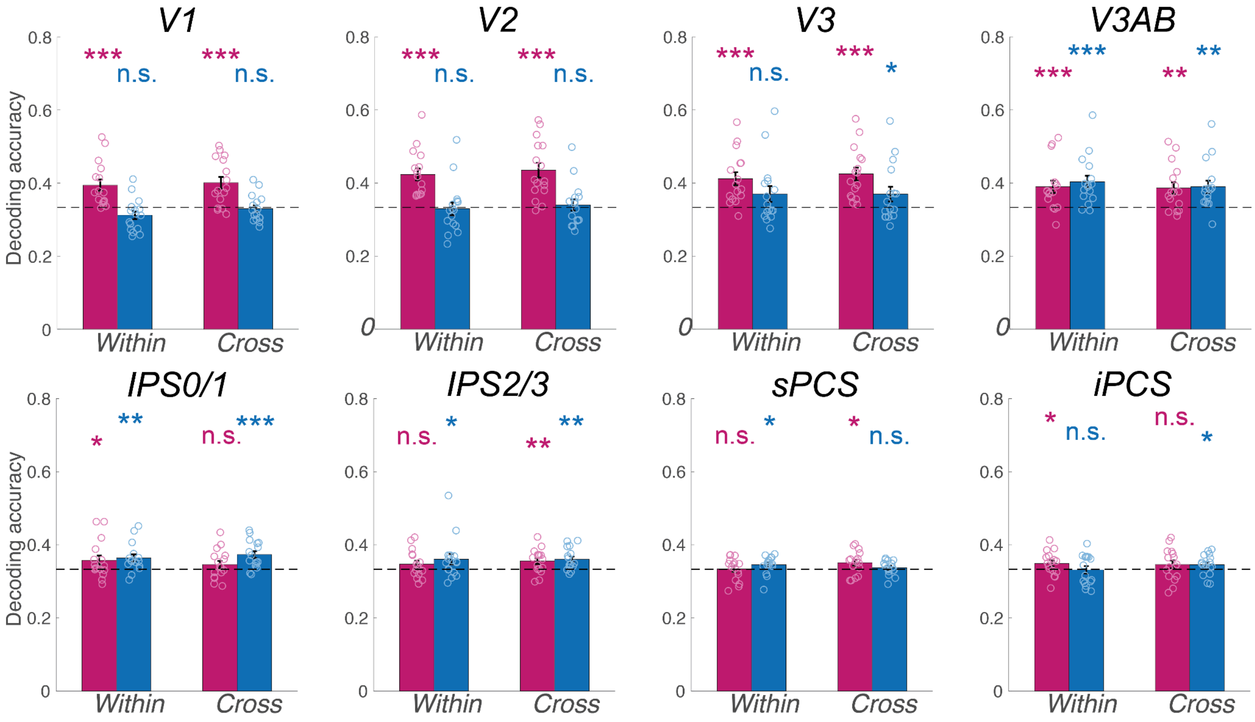


**Figure S3.** Within and cross-modulator decoding results by training classifiers based on the stimulus period of the control task and testing them on the late delay period of the WM task. *$p$<.05, **$p$<.01, ***$p$<.001, n.s. Not significant.

Error bars represent ±1 SEM. Small circles for each bar represent individual data. Dashed horizontal line denotes theoretical chance level (1/3), but results are based on non-parametric permutation tests.

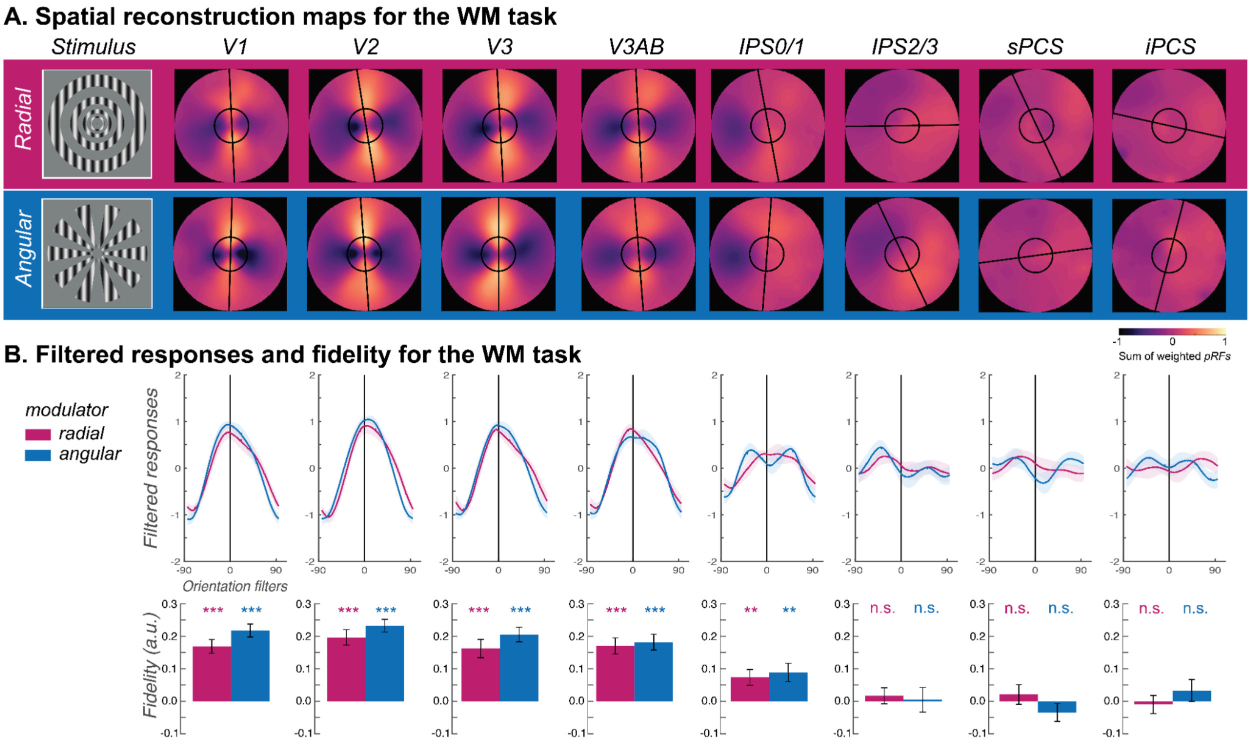


**Figure S4**. Spatial reconstruction results for the WM task across all ROIs
**A.** Line-like patterns emerged across maps of visual space matching the memorized orientation of carrier gratings regardless of the type of modulator (radial - magenta; angular - blue) during the late delay period of the WM task. Spatial maps were rotated such that all orientations were aligned at 0° (top). The warmer colors correspond to increased amplitude of BOLD activity in voxels with receptive fields corresponding to that portion of the visual field. Best fitting lines (black lines) and the size of the stimulus (black circles) are overlaid. **B.** Filtered responses (top row) represent the sum of pixel values within the area of a line-shaped mask (12° length) oriented -90° to 90°, where 0° represents the true orientation. Fidelity values (bottom row) are the result of projecting the filtered responses to 0° (see Methods), where higher fidelity values indicate stronger stimulus orientation representations.

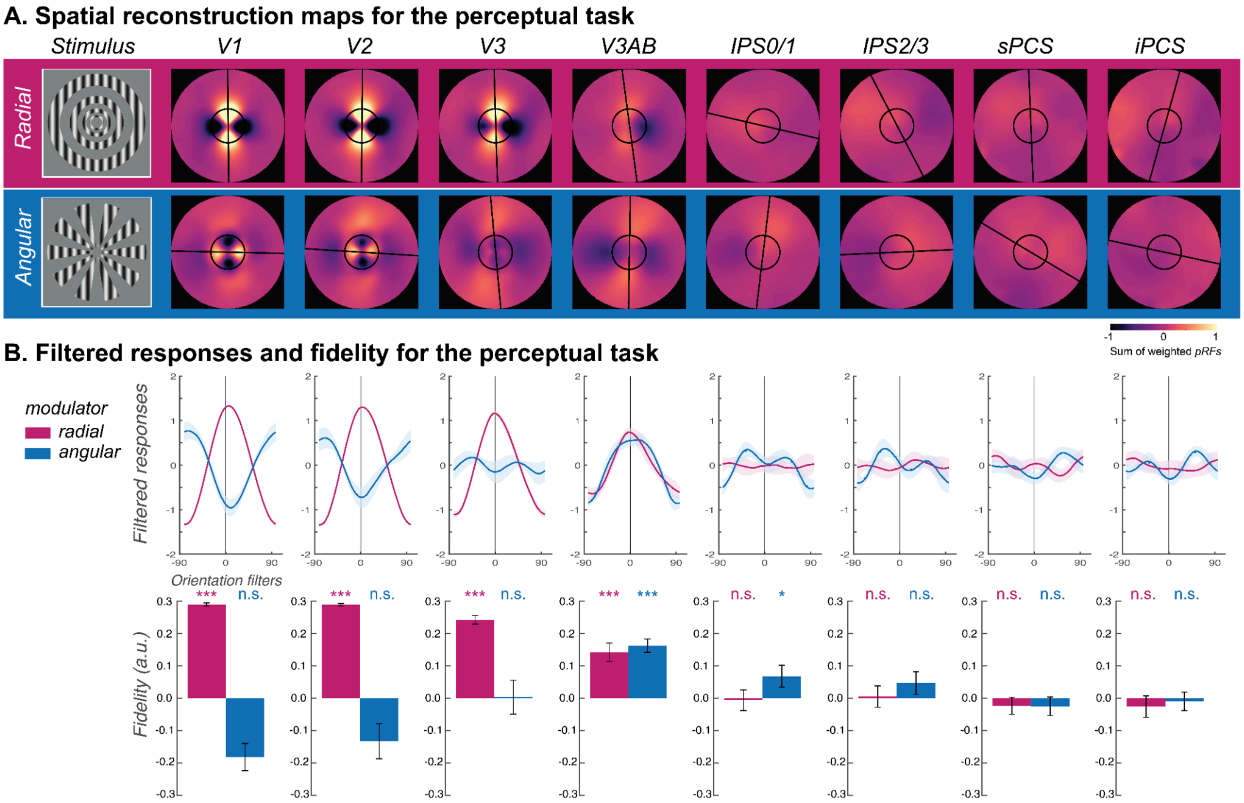


**Figure S5**. Spatial reconstruction results for the perceptual control task across all ROIs

**A.** Line-like patterns emerged across maps of visual space matching the memorized orientation of carrier gratings regardless of the type of modulator (radial - magenta; angular - blue) during the delay period of the perceptual task. Spatial maps were rotated such that all orientations were aligned at 0° (top). The warmer colors correspond to increased amplitude of BOLD activity in voxels with receptive fields corresponding to that portion of the visual field. Best fitting lines (black lines) and the size of the stimulus (black circles) are overlaid. **B.** Filtered responses (top row) represent the sum of pixel values within the area of a line-shaped mask (12° length) oriented -90° to 90°, where 0° represents the true orientation. Fidelity values (bottom row) are the result of projecting the filtered responses to 0° (see Methods), where higher fidelity values indicate stronger stimulus orientation representations.

### Supplementary Table

Table S1.

Decoding accuracy for the stimulus-presenting epoch in the perceptual control task

| One-sample t-test against chance (1/3) based on non-parametric permutation test | | | | | | | | |
|---|---|---|---|---|---|---|---|---|
| | V1 | V2 | V3 | V3AB | IPS0/1 | IPS2/3 | sPCS | iPCS |
| Within radial | **$t$ = 10.433 $p$ < .001** | **$t$ = 15.598 $p$ < .001** | **$t$ = 9.802 $p$ < .001** | **$t$ = 3.848 $p$ < .001** | **$t$ = 2.049 $p$ = 0.033** | $t$ = 0.606 $p$ = 0.245 | $t$ = 0.712 $p$ = 0.259 | $t$ = 0.039 $p$ = 0.510 |
| Within angular | **$t$ = 8.885 $p$ < .001** | **$t$ = 6.339 $p$ < .001** | **$t$ = 7.416 $p$ < .001** | **$t$ = 3.879 $p$ < .001** | $t$ = 1.532 $p$ = 0.067 | **$t$ = 4.691 $p$ < .001** | $t$ = -0.218 $p$ = 0.594 | $t$ = 0.914 $p$ = 0.172 |
| Cross radial to angular | $t$ = -5.255 $p$ = 1.000 | $t$ = -2.993 $p$ = 0.994 | $t$ = -0.406 $p$ = 0.621 | $t$ = 1.669 $p$ = 0.056 | $t$ = 1.029 $p$ = 0.153 | **$t$ = 1.702 $p$ = 0.049** | $t$ = 1.535 $p$ = 0.074 | $t$ = -1.155 $p$ = 0.873 |
| Cross angular to radial | $t$ = -5.660 $p$ = 1.000 | $t$ = -3.342 $p$ = 0.998 | $t$ = -0.598 $p$ = 0.692 | **$t$ = 2.912 $p$ = 0.008** | **$t$ = 1.880 $p$ = 0.040** | **$t$ = 2.483 $p$ = 0.009** | $t$ = 0.470 $p$ = 0.340 | $t$ = -0.751 $p$ = 0.787 |
| Permutation-based 2-way ANOVA. Decoding type × Modulator type | | | | | | | | |
| | V1 | V2 | V3 | V3AB | IPS0/1 | IPS2/3 | sPCS | iPCS |
| Decoding type df: (1, 15) | **$F$ = 233.710 $p$ < .001** | **$F$ = 170.450 $p$ < .001** | **$F$ = 68.036 $p$ < .001** | **$F$ = 7.515 $p$ = 0.007** | $F$ = 1.145 $p$ = 0.290 | $F$ = 1.875 $p$ = 0.179 | $F$ = 0.447 $p$ = 0.520 | $F$ = 1.659 $p$ = 0.202 |
| Modulator type df: (1, 15) | **$F$ = 6.676 $p$ = 0.018** | **$F$ = 12.017 $p$ = 0 .001** | **$F$ = 4.624 $p$ = 0.037** | $F$ < .001 $p$ = 0.993 | $F$ = 0.324 $p$ = 0.576 | **$F$ = 7.597 $p$ = 0.005** | $F$ = 0.778 $p$ = 0.375 | $F$ = 0.428 $p$ = 0.510 |
| Decoding type × Modulator type df: (1, 15) | $F$ = 1.251 $p$ = 0.274 | **$F$ = 7.585 $p$ = 0.009** | $F$ = 3.887 $p$ = 0.051 | $F$ = 0.632 $p$ = 0.435 | $F$ = 0.243 $p$ = 0.621 | $F$ = 2.940 $p$ = 0.094 | $F$ = 0.003 $p$ = 0.959 | $F$ = 0.035 $p$ = 0.855 |

Table S2.

Decoding accuracy for the late delay epoch in the WM task

| One-sample t-test against chance (1/3) based on non-parametric permutation test | | | | | | | | |
|---|---|---|---|---|---|---|---|---|
| | V1 | V2 | V3 | V3AB | IPS0/1 | IPS2/3 | sPCS | iPCS |
| Within radial | **$t$ = 5.302 $p$ < .001** | **$t$ = 4.520 $p$ < .001** | **$t$ = 4.337 $p$ < .001** | **$t$ = 5.891 $p$ < .001** | **$t$ = 4.910 $p$ < .001** | **$t$ = 5.141 $p$ < .001** | **$t$ = 2.706 $p$ = 0.007** | **$t$ = 2.164 $p$ = 0.025** |
| Within angular | **$t$ = 5.279 $p$ < .001** | **$t$ = 6.040 $p$ < .001** | **$t$ = 4.737 $p$ < .001** | **$t$ = 6.260 $p$ < .001** | **$t$ = 5.225 $p$ < .001** | **$t$ = 3.460 $p$ < .001** | **$t$ = 2.936 $p$ = 0.008** | $t$ = 1.718 $p$ = 0.058 |
| Cross radial to angular | **$t$ = 4.513 $p$ < .001** | **$t$ = 4.400 $p$ < .001** | **$t$ = 4.015 $p$ < .001** | **$t$ = 4.953 $p$ < .001** | **$t$ = 4.102 $p$ < .001** | **$t$ = 3.755 $p$ < .001** | **$t$ = 1.835 $p$ = 0.049** | $t$ = 0.476 $p$ = 0.354 |
| Cross angular to radial | **$t$ = 4.695 $p$ < .001** | **$t$ = 4.361 $p$ < .001** | **$t$ = 3.993 $p$ < .001** | **$t$ = 5.239 $p$ < .001** | **$t$ = 4.764 $p$ < .001** | **$t$ = 3.873 $p$ < .001** | **$t$ = 4.636 $p$ < .001** | **$t$ = 2.588 $p$ = 0.009** |
| Permutation-based 2-way ANOVA. Decoding type × Modulator type | | | | | | | | |
| | V1 | V2 | V3 | V3AB | IPS0/1 | IPS2/3 | sPCS | iPCS |
| Decoding type df: (1, 15) | $F$ = 2.553 $p$ = 0.119 | $F$ = 0.931 $p$ = 0.339 | $F$ = 0.232 $p$ = 0.646 | $F$ = 0.426 $p$ = 0.527 | $F$ = 0.138 $p$ = 0.721 | $F$ = 0.012 $p$ = 0.912 | $F$ = 1.172 $p$ = 0.290 | $F$ = 0.652 $p$ = 0.415 |
| Modulator type df: (1, 15) | $F$ = 0.777 $p$ = 0.378 | $F$ = 0.153 $p$ = 0.699 | $F$ = 0.205 $p$ = 0.655 | $F$ = 0.008 $p$ = 0.926 | $F$ = 0.017 $p$ = 0.900 | $F$ = 0.003 $p$ = 0.956 | $F$ = 0.624 $p$ = 0.433 | $F$ = 1.017 $p$ = 0.325 |
| Decoding type × Modulator type df: (1, 15) | $F$ = 1.173 $p$ = 0.283 | $F$ = 0.448 $p$ = 0.509 | $F$ = 0.161 $p$ = 0.684 | $F$ = 0.024 $p$ = 0.876 | $F$ = 0.240 $p$ = 0.624 | $F$ = 0.207 $p$ = 0.656 | $F$ = 0.092 $p$ = 0.771 | $F$ = 0.255 $p$ = 0.613 |

Table S3.

Decoding accuracy for the cross-task decoding by training the classifier in the perceptual control task and testing it in the WM task

| One-sample t-test against chance (1/3) based on non-parametric permutation test | | | | | | | | |
|---|---|---|---|---|---|---|---|---|
| | V1 | V2 | V3 | V3AB | IPS0/1 | IPS2/3 | sPCS | iPCS |
| Within radial | ***t* = 3.961 *p* < .001** | ***t* = 5.596 *p* < .001** | ***t* = 4.430 *p* < .001** | ***t* = 3.373 *p* < .001** | ***t* = 1.927 *p* = 0.031** | *t* = 1.462 *p* = 0.082 | *t* = 0.080 *p* = 0.482 | ***t* = 1.913 *p* = 0.042** |
| Within angular | *t* = -1.907 *p* = 0.963 | *t* = -0.213 *p* = 0.598 | *t* = 1.711 *p* = 0.050 | ***t* = 4.187 *p* < .001** | ***t* = 3.143 *p* = 0.002** | ***t* = 1.923 *p* = 0.036** | ***t* = 2.010 *p* = 0.019** | *t* = -0.183 *p* = 0.574 |
| Cross radial to angular | ***t* = 4.234 *p* < .001** | ***t* = 5.137 *p* < .001** | ***t* = 5.223 *p* < .001** | ***t* = 3.378 *p* = 0.004** | *t* = 1.282 *p* = 0.106 | ***t* = 2.752 *p* = 0.005** | ***t* = 2.215 *p* = 0.014** | *t* = 1.207 *p* = 0.132 |
| Cross angular to radial | *t* = -0.395 *p* = 0.632 | *t* = 0.444 *p* = 0.336 | ***t* = 1.821 *p* = 0.042** | ***t* = 3.410 *p* = 0.003** | ***t* = 4.227 *p* < .001** | ***t* = 3.652 *p* = 0.002** | *t* = 0.798 *p* = 0.212 | ***t* = 1.717 *p* = 0.049** |
| Permutation-based 2-way ANOVA. Decoding type × Modulator type | | | | | | | | |
| | V1 | V2 | V3 | V3AB | IPS0/1 | IPS2/3 | sPCS | iPCS |
| Decoding type df: (1, 15) | *F* = 0.848 *p* = 0.371 | *F* = 0.403 *p* = 0.523 | *F* = 0.105 *p* = 0.749 | *F* = 0.273 *p* = 0.598 | *F* = 0.011 *p* = 0.916 | *F* = 0.176 *p* = 0.691 | *F* = 0.410 *p* = 0.511 | *F* = 0.378 *p* = 0.539 |
| Modulator type df: (1, 15) | ***F* = 33.587 *p* < .001** | ***F* = 30.100 *p* < .001** | ***F* = 6.287 *p* = 0.016** | *F* = 0.260 *p* = 0.612 | *F* = 2.488 *p* = 0.122 | *F* = 0.810 *p* = 0.380 | *F* = 0.026 *p* = 0.879 | *F* = 1.009 *p* = 0.312 |
| Decoding type × Modulator type df: (1, 15) | *F* = 0.167 *p* = 0.690 | *F* < .001 *p* = 0.975 | *F* = 0.127 *p* = 0.728 | *F* = 0.090 *p* = 0.772 | *F* = 0.993 *p* = 0.329 | *F* = 0.198 *p* = 0.660 | *F* = 3.517 *p* = 0.063 | *F* = 0.825 *p* = 0.361 |

Table S4.

Reconstruction fidelity values for the late delay epoch in the WM task

| One-sample t-test against zero based on non-parametric permutation test | | | | | | | | |
|---|---|---|---|---|---|---|---|---|
| | V1 | V2 | V3 | V3AB | IPS0/1 | IPS2/3 | sPCS | iPCS |
| Radial | ***t* = 8.061 *p* < .001** | ***t* = 8.233 *p* < .001** | ***t* = 5.825 *p* < .001** | ***t* = 6.891 *p* < .001** | ***t* = 3.025 *p* = 0.004** | *t* = 0.672 *p* = 0.263 | *t* = 0.686 *p* = 0.240 | *t* = -0.359 *p* = 0.642 |
| Angular | ***t* = 10.875 *p* < .001** | ***t* = 11.998 *p* < .001** | ***t* = 9.070 *p* < .001** | ***t* = 7.559 *p* < .001** | ***t* = 3.132 *p* = 0.004** | *t* = 0.113 *p* = 0.492 | *t* = -1.247 *p* = 0.870 | *t* = 0.975 *p* = 0.149 |

Table S5.

Reconstruction fidelity values for the stimulus-presenting epoch in the perceptual control task

| One-sample t-test against zero based on non-parametric permutation test | | | | | | | | |
|---|---|---|---|---|---|---|---|---|
| | V1 | V2 | V3 | V3AB | IPS0/1 | IPS2/3 | sPCS | iPCS |
| Radial | ***t* = 57.306 *p* < .001** | ***t* = 72.482 *p* < .001** | ***t* = 18.235 *p* < .001** | ***t* = 5.002 *p* < .001** | *t* = -0.183 *p* = 0.562 | *t* = 0.157 *p* = 0.436 | *t* = -0.886 *p* = 0.813 | *t* = -0.781 *p* = 0.788 |
| Angular | *t* = -4.339 *p* = 1.000 | *t* = -2.451 *p* = 0.993 | *t* = 0.059 *p* = 0.467 | ***t* = 7.945 *p* < .001** | ***t* = 2.013 *p* = 0.037** | *t* = 1.351 *p* = 0.110 | *t* = -0.864 *p* = 0.818 | *t* = -0.338 *p* = 0.644 |
